# A tentacle for every occasion: comparing the hunting tentacles and sweeper tentacles, used for territorial competition, in the coral *Galaxea fascicularis*

**DOI:** 10.1101/2020.02.05.936005

**Authors:** Oshra Yosef, Yotam Popovits, Assaf Malik, Maya Ofek-Lalzer, Tali Mass, Daniel Sher

## Abstract

Coral reefs are among the most diverse, complex and densely populated marine ecosystems. To survive, morphologically simple and sessile cnidarians have developed mechanisms to catch prey, deter predators and compete with adjacent corals for space, yet the mechanisms underlying these functions are largely unknown. Here, we characterize the histology, toxic activity and gene expression patterns in two different types of tentacles from the scleractinian coral *Galaxea fascilcularis* – catch tentacles (CTs), used to catch prey and deter predators, and sweeper tentacles (STs), specialized tentacles used for territorial aggression. STs exhibit more mucocytes and higher expression of mucin genes than CTs, and lack the ectodermal cilia used to deliver food to the mouth and remove debris. STs and CTs also express different sensory g-protein coupled receptors, suggesting they may employ different sensory pathways. Each tentacle type has a different complement of stinging cells (nematocytes), and the expression in the two tentacles of genes encoding structural nematocyte proteins suggests the stinging cells develop within the tentacles. CTs have higher neurotoxic and hemolytic activities, consistent with a role in prey capture, whereas the STs have higher phospholipase A2 activity, which we speculate may have a role in inducing tissue damage during territorial aggression. The toxin genes expressed in each tentacle are also different. These results show that the same organism utilizes two distinct tentacle types, each equipped with a different venom apparatus and toxin composition, for prey capture and defense and for territorial aggression.

## Introduction

Competition for space among sessile organisms is a common process in the coral reef environment. The constant battle for space, where each organism aims to obtain the optimal access to light, nutrients and food, has resulted in the development of diverse competitive strategies among reef organisms, including the scleractinian corals that form the physical structure of the reef (Chadwick & Morrow, 2011; Connell et al., 2004). As a result, territorial aggression between corals strongly affects the community competition and resulting reef structure (Jackson & Buss, 1975).

Scleractinian corals employ several methods for territorial aggression (reviewed by (Chadwick & Morrow, 2011; Lang & Chornesky, 1990; R. B. Williams, 1991), for underwater videos of some of these encounters see (Mullen et al., 2016)). Aggressive encounters can operate over a distance, when mediated by water-borne allelochemicals (Atrigenio, Ali, #xf1, o, & Conaco, 2017; Fearon & Cameron, 1997). At closer range, mucous produced by solitary polyps of the family Fungiidae can cause the degradation of adjacent corals (Chadwick, 1988), whereas overgrowth of slow-growing corals by faster-growing ones, reducing the access to light and food, can also be considered a form of aggression (Chadwick & Morrow, 2011). Finally, when two corals come into direct physical contact, they may attack each other using their mesenterial filaments (which are usually used for digestion, (Lang, 1973)), “sweeper polyps” (Peach & Hoegh-Guldberg, 1999) or specialized aggression appendages termed sweeper tentacles, which are the focus of the current study.

Sweeper tentacles (STs) develop on the periphery of coral colonies belonging to several different scleractinian families (R. B. Williams, 1991), and have been observed also in black corals (Goldberg, Grange, Taylor, & Zuniga, 1990) and gorgonian sea fans (Sebens & Miles, 1988). In the scleractinian coral *Galaxea fascicularis*, STs likely develop from the catch tentacles used for feeding (CTs), since intermediate stages, presenting the features of both catch and sweeper tentacles, can sometimes be observed (Hidaka & Yamazato, 1984). The morphogenesis of STs occurs over several days to weeks following the initiation of contact with competitors (or damage caused by competition), although they sometimes develop without any clear cue (Chornesky, 1983; Einat & Nanette, 2006). At the end of their development, STs can be up to 30 times longer than CTs (Figure 1B), and have not been observed to participate in feeding. The contact of a ST with the tissue of a target organism results in tissue damage (lesions) which can also be associated with a cessation of skeleton growth along the contact region between the two competing colonies (Figure 1A). In at least some organisms, STs are temporary structures, regressing after destroying the opponent’s tissue (Chornesky, 1983).

**Figure 1:**
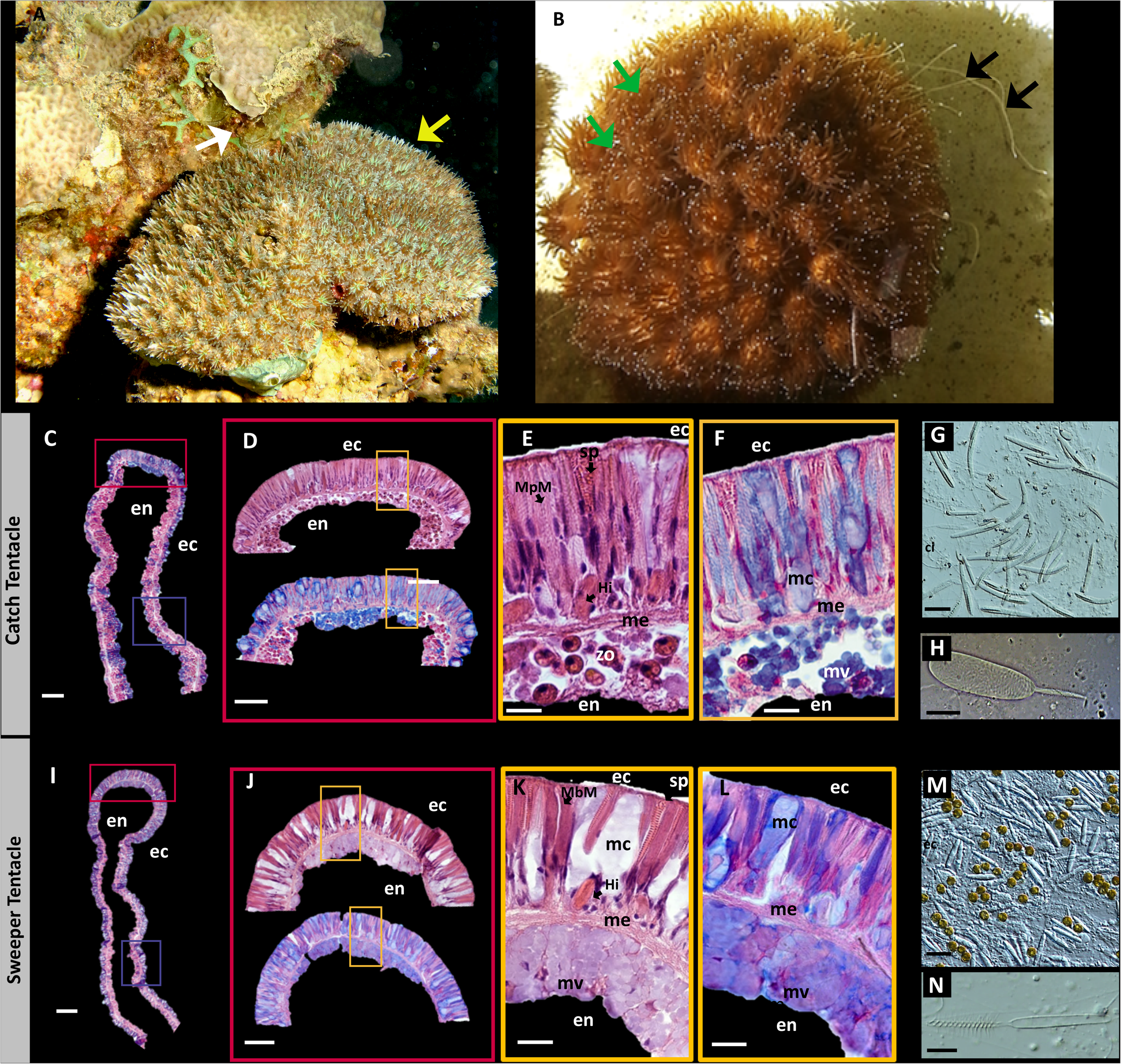
Overview of *G. fascicularis* and its two tentacle types. A) The result of aggressive behavior between Galaxea fascicularis and Pavona sp. The yellow arrow shows the Galaxea, the white arrow shows the dead area between the two corals. Photo taken in Eilat by Tali Mass. B) Catch tentacles (CT, green arrow) and extended sweeper tentacles (ST, black arrow) of G. fascicularis. C and I) Overall histology of Alcian Blue-stained CT (C) and ST (I). Red boxes show the approximate location of high-magnification micrographs in this figure, blue squares show approximate location of micrographs in figure 2. D-F) Histological sections of the tip of the CT, stained with H&E (upper section in D and panel E) and with Alcian Blue (lower section in D and panel F). E and F are magnifications of the orange square in D. Note the tightly-packed large nematocytes in the ectoderm (ec) and the abundance of symbiotic algae (zo) in the endoderm (en). G, H) Main types of nematocytes of the acrosphere (Microbasic p Mastigophores, MbMs, which have a smooth shaft, unlike the barbed shaft of the MbMs of the sweeper tentacles). I-L) Histological sections of the tip of the ST, stained with H&E (upper section in J and panel K) and with Alcian Blue (lower section in J and panel L). K and L are magnifications of the yellow square in J. Note the mucocytes in the ectoderm between the large, elongated nematocytes, and the lack of symbionts in the endoderm. Mucocytes (mc), mucus vesicles (mv) and nematocysts are stained in blue, the latter perhaps due to the presence of poly-gamma-glutamate (an acidic polyanion) in the capsule matrix of the nematocysts. M, N) Main types of nematocytes of the acrosphere of the ST (Microbasic b Mastogophores, MbMs). Scale bars are 100μm for C and I, 50μm for D and J, 200μm for E, F, K and L, 25μm for G, M and 10μm for H, N.

While the presence of STs in different corals and their ecological role in territorial aggression have been extensively studied (primarily during the 1980’s), little is known about the mechanism of aggression at the molecular and biochemical levels. Cnidarians are known to produce complex and highly effective toxins that are used to catch prey and defend themselves against predators (recently reviewed by (Jouiaei et al., 2015; Schmidt, Daly, & Wilson, 2019)). These toxic cocktails are typically injected into prey or predators through specialized stinging cells called nematocytes (Beckmann & Ozbek, 2012; Tardent, 1995), although some toxins are released from epithelial cells or even deposited into oocytes (Columbus-Shenkar et al., 2018; Moran et al., 2011; Daniel Sher, Fishman, Melamed-Book, Zhang, & Zlotkin, 2008). Most cnidarians, including corals, produce several types of nematocytes, and the nematocytes found in the STs are different from those in the CTs. For example, in *Galaxea fascicularis*, the acrospheres (the tentacle tips, which contain numerous nematocytes) of STs consist of approximately 50% Large Microbasic b-Mastigophores (L-MbM, which penetrate into target tissues and presumably inject toxins) and almost 40% spirocysts (Hidaka & Yamazato, 1984), which do not penetrate into the target tissue and have primarily an adhesive role. In contrast, CTs consist of 80% spirocysts and the remainder are mainly Microbasic p-Mastigophores (MpM), another type of penetrating nematocyte (Hidaka & Yamazato, 1984). Thus, while it is likely that the nematocysts and the toxins they produce have an important role in aggression, the specific mechanisms underlying the aggression at the molecular level are currently unknown.

To obtain insights into the mechanism of territorial aggression, we performed in-depth comparison of the CTs and STs of the Indo-Pacific scleractinian coral *Galaxea fascicularis* at the histological, biochemical and molecular levels. *Galaxea fascicularis* is a massive (boulder-shaped) coral that is known to be highly aggressive, both towards other species and towards genetically-different clades or morphotypes from the same species (Abelson & Loya, 1999; Genin, Karp, & Miroz, 1994). *G. fasciularis* develop ST in response to the presence of a potential competitor (Hidaka & Yamazato, 1984), and may also use their mesenterial filaments for aggression (Evensen & Edmunds, 2018). The STs of *G. fasciularis* have been studied from an organismal-behavioral point of view (*e.g*. the effect of genetic similarity and of flow (Genin et al., 1994; Hidaka, 1985)), and the dynamics of their development have been characterized (Hidaka & Yamazato, 1984). We therefore employed a comparative transcriptomics approach, and we discuss specific genes and pathways differentially expressed between the two tentacle types in relation to detailed observations of the histology of the two tentacle types, their nematocyst complements and their toxicity (neurotoxic, hemolytic and phospholipase A2 activities).

## Materials and Methods

### Specimen collection and histology

tentacle samples were obtained from the underwater marine observatory in Eilat. Paired catch and sweeper tentacles were collected from *G. fascicularis* colonies exhibiting the same morphotype (brown with a green oral disc), all located in the same large aquarium, under the same, uniform abiotic conditions, with water in the aquarium drawn directly from the Red Sea at water depth of 20m. Tentacles for RNA extraction and bioassays were cut from the colonies by a SCUBA diver, placed in 50ml Falcon tubes and immediately brought up to the surface. Excess water was discarded, TRIzol® Reagent was added to the samples used for RNA extraction, the samples were flash-frozen in liquid nitrogen and were maintained at −80°C till further processing. For histological and microscopic analyses, tentacles were either anesthetized in 0.3M magnesium chloride for 10 min, and either observed fresh (for “squash assays”) or fixed in 2% forma-glutaraldehyde solution for 72 hours at 4°C and transferred to 70% ethanol at 4°C till histological processing was performed. Embedding into paraffin blocks, sectioning into 4-micrometer sections and histological staining were all performed using the Ventana BenchMark Fully Automated Slide Stainer System at the Histology Service Center at Carmel Hospital, Haifa Israel. Two staining techniques were used: H&E (hematoxylin and eosin) for cell visualization and Alcian blue for identifying acid mucopolysaccharides and acidic mucins. Microscopic observations were performed using a Nikon Eclipse Ti microscope, with nematocysts identified as previously described (Östman, 2000).

### RNA extraction, transcriptome sequencing and quality control

The tentacles for RNA extraction were collected at 17:00 on March 20^th^, 2017. Samples from the polyp body were also collected for this analysis, in order to allow the assembly of a comprehensive transcriptome database. Total RNA was extracted after thawing the tissues and electric homogenization using TRIzol® followed by purification with the Zymo Quick-RNA™ MiniPrep kit. Libraries were prepared using the Genomics in house protocol at the Weizmann Institute of Sciences Genome Center for mRNA-seq. Briefly, the polyA fraction (mRNA) was enriched from 500 ng of total RNA followed by fragmentation and the generation of double-stranded cDNA. Then, end repair, A base addition, adapter ligation and PCR amplification steps were performed. Libraries were evaluated by Qubit (Thermo fisher scientific) and TapeStation (Agilent). Sequencing libraries were constructed with unique barcodes for each sample to allow multiplexing. Around 18-24 million 125-bp Paired-End reads were sequenced per sample on an Illumina HiSeq High Output instrument. Paired End (PE) reads were adapter-trimmed using cutadapt 1.15 (https://cutadapt.readthedocs.io), low-quality regions were removed with Trimmomatic 3.0 (Bolger, Lohse, & Usadel, 2014) the resulting trimmed reads were inspected using Fastqc (http://www.bioinformatics.babraham.ac.uk). Reads mapping to human genes, NCBI univec databases, ribosomal RNA databases (Bengtsson-Palme et al., 2015), and *Symbiodiniaceae* genes, were filtered out, using Fastqscreen (http://www.bioinformatics.babraham.ac.uk). Since Fastqscreen uses Bowtie2 mapper, which is only designed to align highly similar sequences, not all *Symbiodiniaceae* species could be excluded at this step. Therefore, additional detection methods were used to exclude symbionts after assembly (see below).

### Transcriptome assembly and annotations

The reads after quality control were assembled using Trinity 2.4 (Grabherr et al., 2011) with default settings, except for the addition of Slurm cluster management setting (Trinity arguments: grid_exec, grid_conf). For assembly diagnostics, PE reads were mapped back to the assembly using Bowtie2 default PE settings. 78%-85% of all the PE reads were concordantly aligned to the same contig, which are typical results for such de-novo assembly using Trinity. Trinity produces contigs (equivalent to transcripts), which can be clustered into Trinity-genes. For Trinity genes with open reading frames (ORFs) ≥100aa long (github.com/TransDecoder/TransDecoder), the largest ORF was compared to the NCBI nr database using Blastx. In addition, we used the tools Trinotate (https://trinotate.github.io) and Blast2GO (https://www.blast2go.com) for functional annotations.

### Transcript quantitation and differential expression analysis

PE reads abundance, at the Trinity gene level, was estimated with RSEM (B. Li & Dewey, 2011). From all Trinity genes quantified by RSEM, only genes with unambiguous metazoan-origin were selected (excluding NCBI nr proteins mapped to *Symbiodiniaceae* or other non-metazoa genes, as well as un-annotated genes). Species origin was estimated by first finding the best NCBI NR database hits for the longest ORFs using blastp, and then detecting species origin based on the NCBI gi id to taxonomy database (ftp://ftp.ncbi.nih.gov/pub/taxonomy). Overall, for the different samples, 77%-82% of all quality-filtered reads were mapped with RSEM to 360,065 Trinity genes (most of these genes possibly represent non-coding transcripts). After excluding non-metazoan annotations, 26%-48% of the original quality-filtered reads survived (4.2-8.7 million reads remained per sample, median 7.7 million), in 28,588 Trinity genes, considering the catch and sweeper samples. DE analysis was conducted using Bioconductor DEseq2, using only the samples of the two tentacle types, considering two factors in the DESeq2 GLM models: tissue identity (catch, sweeper) and specimen (paired 3,7,8,9 samples) (Love, Huber, & Anders, 2014). We further verified the general horizontal symmetry of the resulting MA plot in DEseq2 in catch vs. sweeper samples, and also tested for comparable normalized count frequency distribution between samples. NMDS (non-metric multidimensional scaling) ordination by groups was conducted in R using the Vegan package (cran.r-project.org/web/packages/vegan). Gene Ontology annotation and enrichment analyses were performed using BLAST2GO (Conesa et al., 2005), with enrichment defined as the GO terms in each tentacle compared with the full transcriptome. The results were then narrowed down using the “reduce to more specific” function, which removes generic functions if their more specific child term is also significant. For the identification of toxin and GPCR genes, a reciprocal best-BLAST-hit (RBBH, (Rachamim et al., 2015)) approach was utilized. Toxins sequences were from (Rachamim et al., 2015)), genes involved in cilia/flagella and G-protein-coupled receptors (GPCRs) were from (Hall et al., 2017) with additional sequences added from other literature sources. These sequences were searched against the Trinity assembly using BLAST, with the results compared to the Swissprot database.

### Bioassays

The tentacles were thawed, 1ml of PBS (Phosphate Buffered Saline) was added, homogenization was performed using an electric homogenizer for 10 seconds, and the homogenate was centrifuged for 5 minutes at 5000 rpm. The supernatant (crude water extract) was used for all bioassays. Paralytic activity was assessed by injection to *Sarcophaga faculata* blowfly larvae ((Zlotkin, Fraenkel, Miranda, & Lissittzky, 1971), average weight 110±20 mg) and a positive result was scored when full immobilization of the larva was observed within 60 seconds. 5-10μl was injected at each time, with PBS serving as a negative control. PD_50_ (paralytic dose - defined as the amount of protein needed to paralyze 50% of the tested animals) was determined for every sample and was calculated according to the estimation method of Reed and Muench (Reed & Muench, 1938). Hemolysis was performed as described in (Ben-Ari, Paz, & Sher, 2018), using O Rh positive human blood Type obtained from the Yoseftal Hospital Blood Bank.. One hemolytic unit (HU50) was defined as the amount of protein sample required to cause 50% hemolysis. Phospholipase A2 (PLA2) activity was measured using the EnzChek^®^ Assay Kit, using honey bee venom PLA2 as a positive control and to produce a standard curve. The kits limit of detection was 0.05 Units/ml, with one PLA2 unit defined as the amount of protein needed to hydrolyze of 1 μmole of L-aphosphatidylcholine to L-a-lysophosphatidylcholine and a fatty acid per minute at pH 8.0 at 37°C. Protein concentration was measured using the BCA protein kit (Pierce BCA), with Bovine Serum Albumin (BSA) used to produce a standard curve ranging from 2 - 0.2 mg/ml protein. Statistical tests for the bioassays were performed using paired samples *t*- test using IBM SPSS Statistics software. All data was normally distributed in normality tests and significance at a 0.05 level was used for all tests. Since no paralytic activity was detected in sweeper tentacles, results could not be statistically analyzed but differences were clear between tentacle types.

## Results

### Differences in the tissue structure between catch and sweeper tentacles

The Catch tentacles (CTs) and Sweeper Tentacles (STs) of *Galaxea fascicularis* have very different macroscopic morphologies, with the ST being up to 30 times longer than CTs (Figure 1B). To identify the tissue structures potentially underlying these morphological differences, we produced histological sections of the two tentacle types using two different stains – the classical Hematoxylin-Eosin (H&E) stain and Alcian Blue, a cationic dye which stains primarily acidic polysaccharides. As shown in Figures 1 and 2, major histological differences are observed between the tentacles. Starting at ectoderm of the tip of the tentacle (the acrosphere), the CT can be characterized by a dense layer of nematocytes, including mostly spirocysts and Microbasic p-Mastigophores (MpMs) (Figure 1 C-H). In contrast, while many nematocytes were observed also at the tip of the STs, these were less dense than in the CT, and there were many more mucous secreting cells compared with the CT (Figure 1J-L). Unlike the CTs, the nematocysts of the ST were mainly Microbasic b-mastigophpores (MbMs, Figure 1M, N), including a type not observed at all in the STs (very large MbMs). The different types of nematocytes are consistent with the observations of Hidaka (Hidaka & Yamazato, 1984). The endoderm of the two tentacles types was also different, with the CTs characterized by many symbiotic algae residing within the endodermal cells (Figure 1 E, Figure 2) and the STs comprising many mucous vesicles, making it difficult to distinguish individual cells (Figure 1 L).

**Figure 2:**
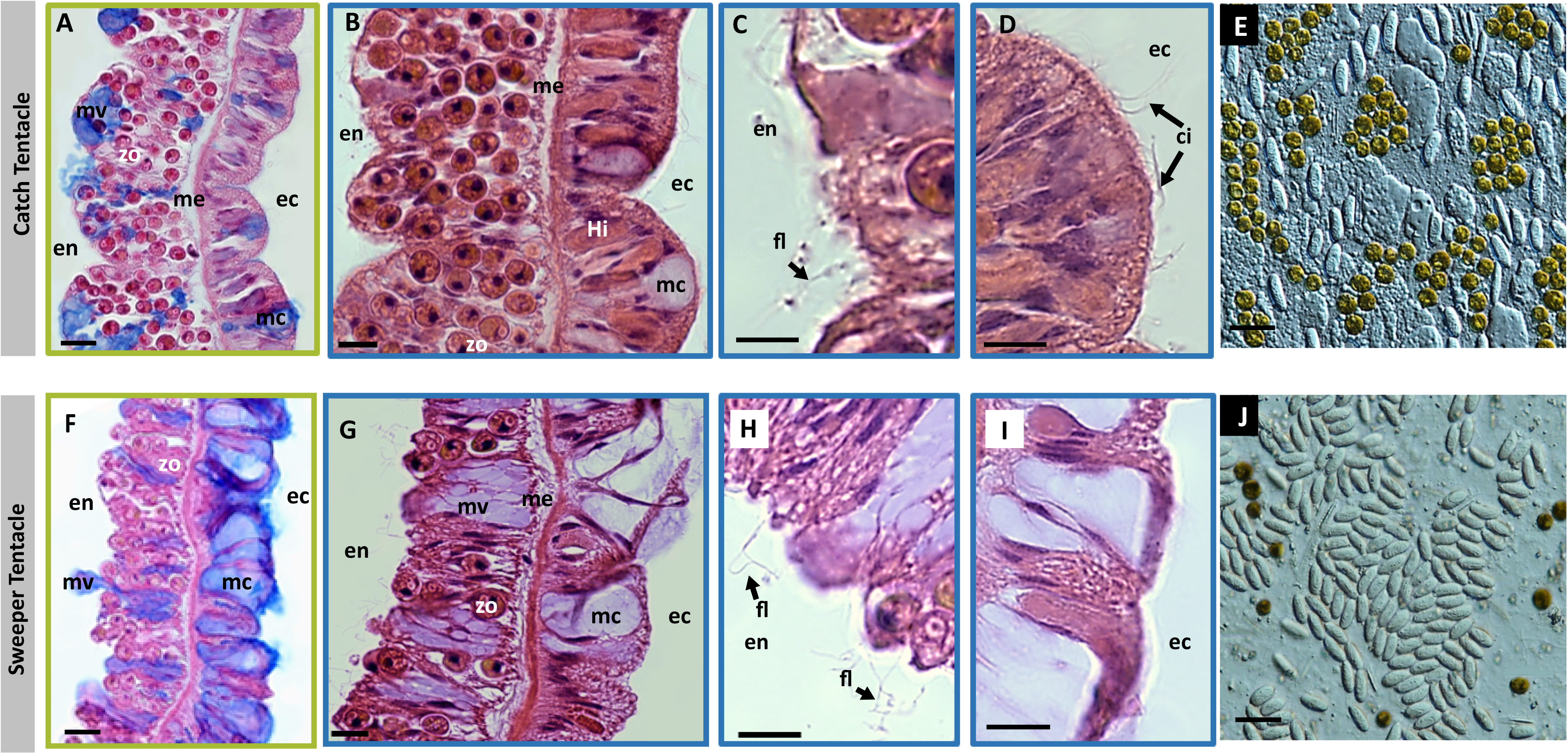
Comparison of the base of the two tentacle types - abundant secretory mucocytes and lack of cilia in the ecotoderm of ST compared to CT. Catch (A-D) and sweeper (E-H) tentacles sections are shown, respectively, stained with Alcian blue (A, F) and H&E (B-D, G-I). The endodermal layer (en) is wider than the ectodermal layer (ec), especially in the CT, with higher number of zooxanthellae symbionts (zo) in the CTs in comparison to the STs (compare A, B with F, G). Flagella (fl) can be observed on the endodermal layer of both tentacle types (C, H), facing the gastrovascular lumen. In contrast, ectodermal cilia (ci) are present only in catch tentacles (D) E, J) Squash assays of the base area of catch and sweeper tentacles respectively, composed mainly from of holotrichous isorhiza nematocysts. Note here too the higher density of zooxanthellae in the CTs. Scale bars: A, B, F, G: 20μm; C, D, H, I: 10μm; E, J 25μm.

Similar to the tips of the tentacles, at the base of the tentacles there was also a much higher density of mucocytes in the STs compared to the CTs (compare Figure 2 A, B to F, G). The endoderm layer of the CTs was wider than that of the STs, and had a higher density of zooxanthellae (Figure 2). The nematocytes at the base of the CTs and STs were similar, comprising mainly holotrichous isorhiza (HI, Figure 2E, J).

A major difference between the CT and ST, observed in Hematoxylin-Eosin stained sections of the two tentacle types, was the lack of ectodermal cilia on the surface of STs (Figure 2 I, compared to D). However, flagella were observed at the endodermal layer of both tentacles type (Figure 2 C, H). Ectodermal cilia are involved in the ciliary-mucoid feeding and cleansing processes, by helping to capture particulate food and transporting it towards the coral’s mouth. They also may help clean the coral surface from sediment entrapped in the mucus (Brown & Bythell, 2005).

### Overview of the transcriptome assembly and differential gene expression patterns

To obtain an overview of the molecular differences between CTs and STs, we sequenced, assembled *de-novo* and annotated transcriptomes from the two tentacle types of four different specimens of *G. fascicularis*. We also sequenced whole-body transcriptomes from three specimens, with the aim of obtaining a comprehensive transcriptome database. The final database comprised a total of 28,588 putative genes, somewhat more than predicted from a recently published draft genome (22,418, (Milde et al., 2009)). Clear differences were observed between the transcriptome profiles of the two tentacle types, and between them and the whole-body samples (Figure 3A). Despite the clear clustering of the samples based on tentacle type, significant variability was observed between the same tentacle types from different colonies (Figure 3B). Such inter-colony variability is in agreement with previous studies (Seneca & Palumbi, 2015). Pairwise comparisons between the two tentacles type revealed that 1585 genes were more highly expressed in the STs compared to 1165 genes more highly expressed the CTs (Supplementary Excel Table). Enrichment analysis showed that 14 Gene Ontology (GO) terms were enriched among the genes more abundantly expressed in the STs, whereas only two were enriched in the CT (Figure 3C). Two terms enriched in the STs, phospholipases and metalloproteases, are suggestive of functions involved in venom toxicity ((Casewell, 2012; Lee et al., 2011), see below). Similarly, the enrichment of GO terms in the STs related to voltage gates calcium channel and ionotropic glutamate receptor activities suggest differences in the pathways of cellular signal transduction and synaptic excitatory transmission between the two tentacle types. Other enriched pathways in the STs include carbohydrate binding, heme and oxygen binding, catalase and several other molecular functions.

**Figure 3:**
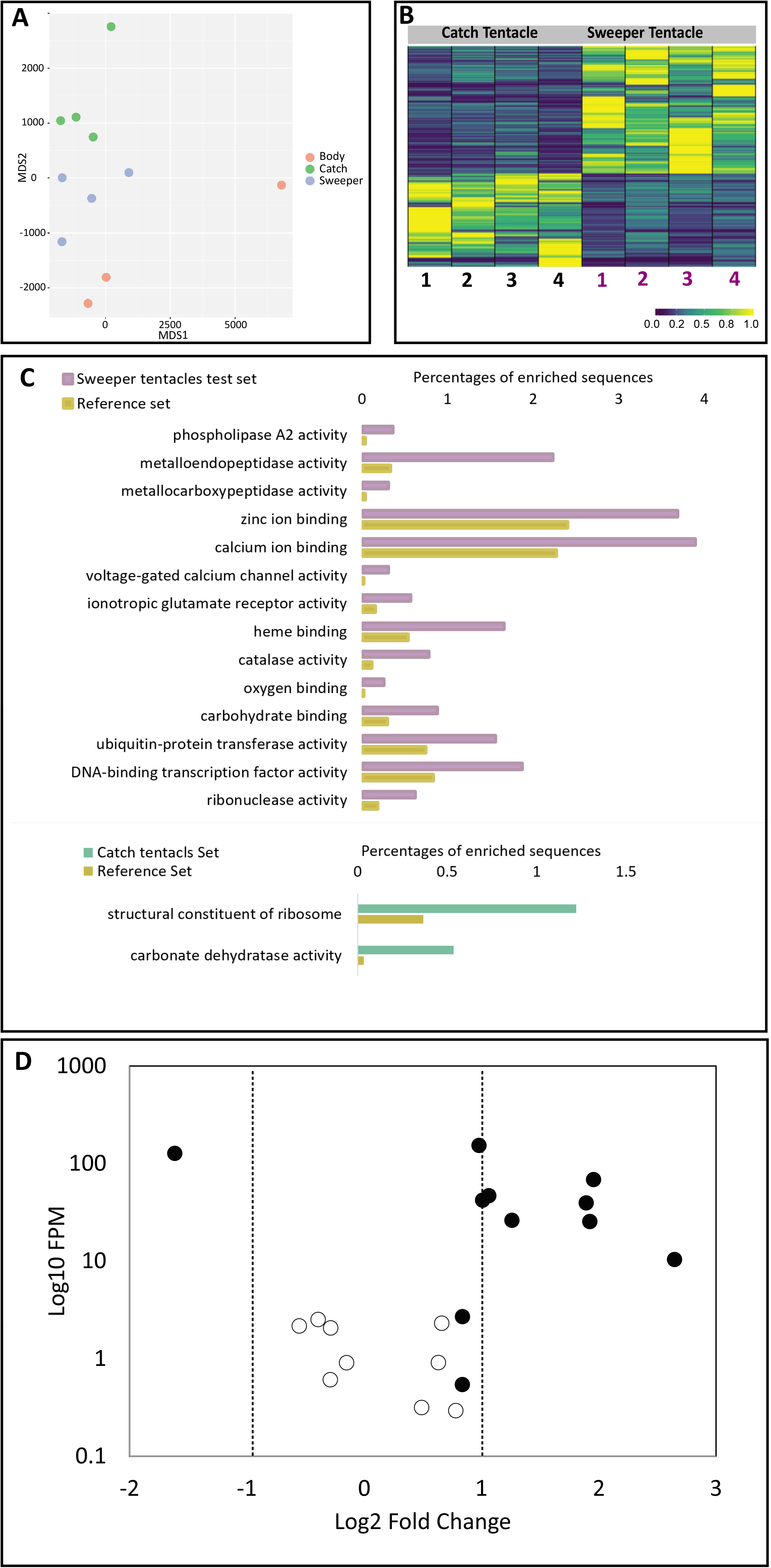
Overview of the gene expression in the two tentacle types. A) Non-metric multidimensional scale plot of Manhattan distance indexes representing three tissue types sweeper tentacles (blue), catch tentacles (green) and whole polyp (red). Each point represents tentacles collected from a single colony of *G. facsicularies*. B) A heatmap showing the expression patterns of differentially expressed genes between the ST and CT (log10 Fragments Per Million or FPM). Note the variability between biological replicates. C) Gene Ontology terms enriched in the sweeper and catch tentacles. The reference represents the abundance of these terms in the full transcriptome. D) Differences in the expression of putative mucin-encoding genes between the CT and the ST. Closed circles represent transcripts whose expression differs significantly between the two tentacle types (adjusted p-value < 0.05).

The broad-scale observations of changes in gene expression led us to examine the expression patterns of specific genes that might be related to differences in the tissue structure of the two tentacle types. Since we observed differences in the distribution of mucocytes, we first asked whether there are differences in the expression of genes encoding mucins. Indeed, eight genes encoding mucins were more abundantly expressed in the STs, compared with only one such gene in the CTs (Figure 3D). In contrast, despite the presence of ectodermal cilia in the CTs but not the STs, no clear differences were observed between the CTs and STs in the expression of genes involved in the synthesis of cilia or flagella (Supplementary Excel file). We also asked whether we could identify differences in the expression of genes encoding nematocyst structural components, which might be related to the different nematocytes found in each tentacle type. While we identified transcripts encoding the nematocyst-specific genes nematogalectin and NOWA, both of which were more abundantly expressed in the STs, no differences in the sequences were observed between the tentacle types (Supplementary Excel File) {Hwang, 2010 #648; Engel, 2002 #649}.

### Sensory G-protein coupled receptor genes are differentially expressed between the CTs and STs

Catch and sweeper tentacles both need to respond to environmental cues, and these cues are likely different – catch tentacles need to rapidly respond to chemical and physical stimuli from motile prey whereas sweeper tentacles exhibit “searching” behavior (Einat & Nanette, 2006), and potentially can respond more slowly as their targets are sessile. Once they have identified their targets, both tentacle types discharge nematocytes. As described above, GO terms related to calcium cellular signaling and ionotropic glutamate receptor activity were enriched in genes more abundantly expressed in the ST (Figure 3C), suggesting differences in the sensory or neuronal circuitry between the tentacle types. Recently, G-protein-coupled receptors (GPCRs) have been implicated in environmental sensing in another marine invertebrate, the Crown-of-thorns sea-star *Acanthaster plancii* (Hall et al., 2017). Motivated by this study, we identified 52 genes encoding rhodopsin-like GPCRs in the full transcriptome data from *G. fascicularis*. Of these genes, 16 and 12 genes found to be more abundantly expressed in the ST and CT respectively, compared to only 6 genes more abundantly expressed in the body tissue (Supplementary Excel file). The GPCR genes more abundantly expressed in the CT included two histamine H2-like receptor genes (out of a total of 3), multiple genes encoding Substance-K receptors and QRFP-like peptide receptors (Supplementary Excel file). In the ST, multiple genes encoding non-visual photoreceptor (NVP) genes such as Melanopsin and Melatonin receptors were more abundantly expressed, as were several other genes encoding receptors for neuropeptides (*e.g*. RYamide and tachykinin). A gene encoding a putative Allostatin receptor, which in *Hydra* was shown to have myoregulatory activity effecting the shape and length of the tentacles (Alzugaray, Hernández-Martínez, & Ronderos, 2016), was also more abundantly expressed in the ST.

### The CTs and STs differ in the expression of toxin-encoding genes in and three types of tissue toxicity

The primary ecological roles of the catch and sweeper tentacles are to affect target organisms, presumably using nematocyst-derived venom or other toxins. The venom of cnidarians has been studied extensively, and is comprised primarily of proteins and polypeptides, including neurotoxins, pore-forming hemolysins, phospholipase A2 (PLA2) toxins and a wide variety of enzymes such as proteases (recently reviewed by (Jouiaei et al., 2015; Schmidt et al., 2019)). Therefore, to begin elucidating the molecular mechanism underlying the different ecological functions of the venoms, we searched for known cnidarian toxins in the transcriptomes of the CTs and STs. In total, we identified 23 genes encoding putative toxins that belong to four different classes of toxins: hemolytic toxins, phospholipase enzymes toxins, metalloprotease and Kunitz type toxins (Figure 4A). We also measured the paralytic, hemolytic and phospholipase A2 activity of tissue extracts from both tentacle types, in order to seek potential relationships between the expressed genes and actual toxic activities.

**Figure 4:**
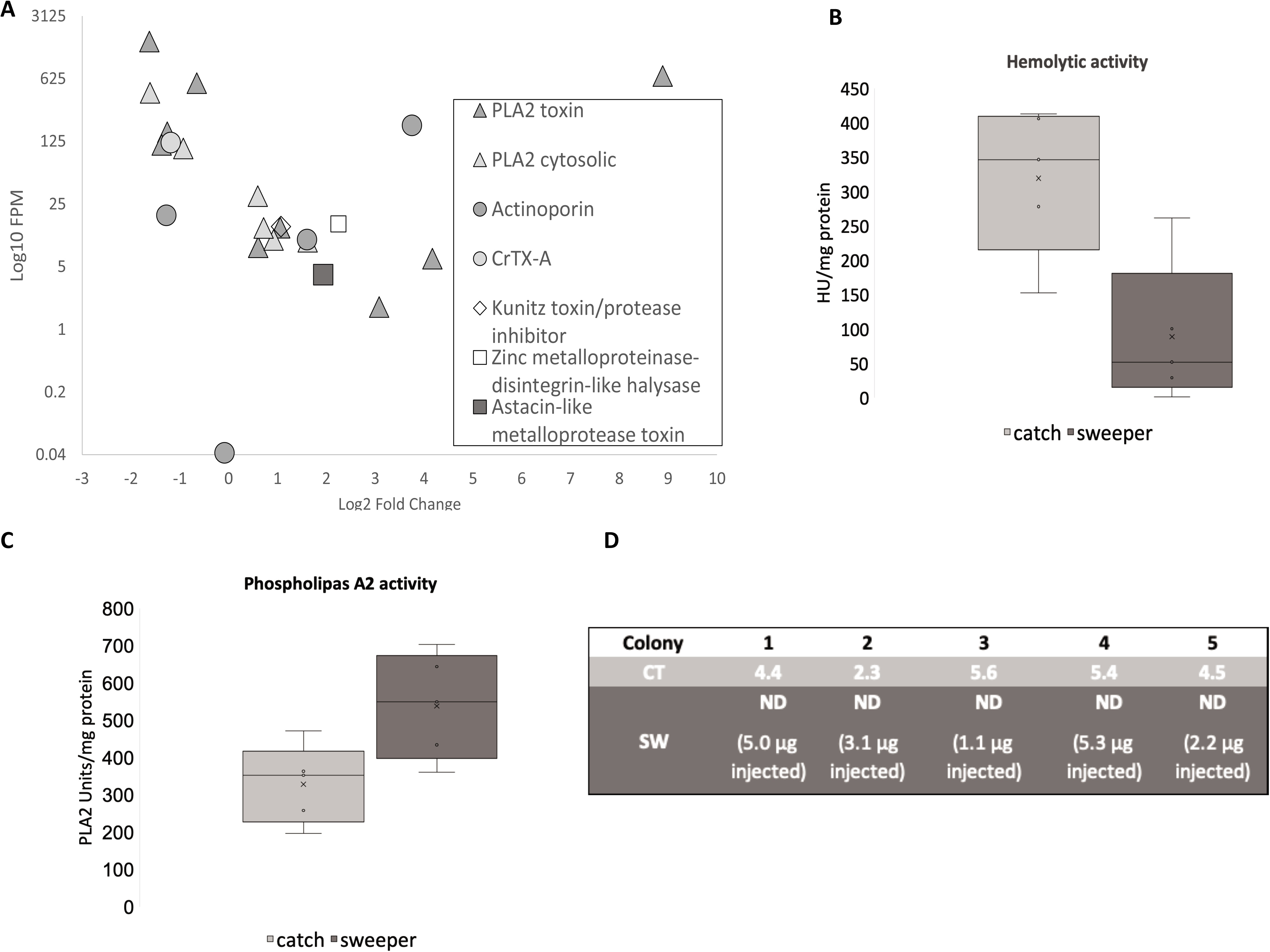
Sweeper and catch tentacles toxic assays and toxins gene expression results. A) Differences in the expression of putative toxin-encoding genes between the CT and the ST. the differences in expression of all of the genes with more than 2-fold difference were statistically significant (adjusted p-value <0.05). B) hemolytic activity is lower in the sweeper tentacles compared to the catch tentacles. Hemolytic units (HU) are shown, normalized to milligram protein. N=5, t(4)= 3.911, p<0.001. C) Phospholipase A2 (PLA2) activity is higher in sweeper tentacles compared to catch tentacles. PLA2 activity is presented as the number of PLA2 units normalized to milligram protein. t(4)=-3.213, p=0.032. Boxes are the interquartile range, lines are medians N=5. D) Paralytic activity was detected in catch tentacles but not in sweeper tentacles. The results shown are the PD_50_ (dose, in micrograms, required to paralyze a ~110mg blowfly larva within 1 minute).

We identified five genes encoding putative hemolytic toxins, belonging to two distinct families: Actinoporins and CrTX-A like toxins (Figure 4A, Supplementary Excel file). A total of four distinct actinoporin genes were identified, with two of the isoforms more abundantly expressed in the STs and one in the CTs (Figure 4A). A single gene encoded a putative hemolytic toxin from a different family, similar to CrTX-A from the Box Jellyfish Carybdea rastonii which is potently hemolytic and lethal by injection mice and crayfish (Nagai et al., 2000). The CrTX-A like toxin was expressed 3-fold more in the CTs compared with the ST (Figure 4A). In agreement with the expression of genes encoding putative hemolysins, both tentacle types exhibited hemolytic activity, with the CT extracts being approximately 3-fold more hemolytic than the STs (Figure 4B).

We also identified eight different genes encoding putative Phospholipase A2 (PLA2) toxins, which had the highest levels of expression and also the highest fold change between tentacle types of all putative toxin-encoding genes (*e.g*. up to ~900-fold higher expression level of one gene in the STs compared with the CTs, Figure 4A). Four of the putative PLA2-encoding genes were more abundantly expressed in the CTs and four in the STs (Figure 4A). In this case too, PLA2 activity was identified in the extracts of both tentacles, but in contrast to the hemolytic activity was significantly stronger in the STs (Figure 4C). Genes encoding metalloproteases were also identified, and were more abundantly expressed in the STs (Figure 4A).

Paralyzing prey organisms is the basic role of cnidarian venom used to catch prey, and, as expected, the CTs exhibited paralytic activity (Figure 4D). No paralysis was observed in any of the larvae injected with the water extract of the STs, even at the highest injected doses (Figure 4D). In addition, all larvae injected with the extract from the CTs died within several hours of injection, whereas all larvae injected with the extracts from the STs survived after 24 hours. We note that due to the limited amount of ST biomass available, in some pairs (*e.g*. from colonies 3 and 5), the injected dose from the STs was below the dose required to paralyze 50% of the injected larvae (PD50) of the catch tentacles, thus precluding a direct statistical test of the difference in paralytic activity. Nevertheless, these results suggest that even if paralytic toxins are found in the STs, their concentration and/or activity is much lower than those of the catch tentacles. We were unable to identify transcripts encoding any of the known peptide toxins targeting Na^+^ or K^+^ channels (Jouiaei et al., 2015), with the exception of a putative Kunitz-type toxin. Such proteins can potentially inhibit K^+^ channels but can also act as a protease inhibitors (Pritchard & Dufton, 1999). However, the expression of the putative Kunitz type toxin was slightly higher in the STs, suggesting that it is not the molecule responsible for the paralytic activity.

## Discussion

Over millions of years, scleractinian corals have evolved methods to aggressively compete for territory on the often-crowded coral reef. These aggressive interactions are one of the main factors leading to the complex physical structure of coral reefs, which provides the substrate and shelter for some of the worlds’ most biodiverse ecosystems (Chadwick & Morrow, 2011; Connell et al., 2004). While a variety of aggressive mechanisms have been described, molecular mechanisms of underpinning these interactions, and how they are related to the morphology and histology of the interacting organisms, are mostly unknown. In the following discussion, we use the transcriptomic results combined with observations based on histology and toxicity bioassays to begin elucidating the mechanisms of interaction in *Galaxea fascicularis*, one of the more well-studied models of coral aggression (Evensen & Edmunds, 2018; Genin et al., 1994; Hidaka, 1985; Hidaka & Yamazato, 1984).

### Histological adaptations for prey capture vs territorial aggression

Several major histological differences were observed between the CTs and the STs, with perhaps the most apparent being the lack of ectodermal cilia and the higher density of mucocytes in the ectoderm of the STs (including at the base of the tentacles, Figure 2). The coral mucus is essential for many aspects of coral biology, including heterotrophic feeding, protection from parasites, pathogens and environmental stresses, sediment cleansing and structural support (Brown & Bythell, 2005). In addition, in cnidarians, mucous may play both modulatory and effector roles in territorial aggression. Mucous may play a role in allorecognition by cnidarians, for example by promoting the discharge of nematocytes against other species while inhibiting the discharge when encountering the same species or other tentacles from the same organism (Ertman & Davenport, 1981; Gundlach & Watson, 2018). It may also have a direct alleloptahic role, causing damage to target organisms (Chadwick, 1988), and may contain bioactive molecules or toxins (Stabili, Schirosi, Parisi, Piraino, & Cammarata, 2015). It is noteworthy that some of the most aggressive corals, including *Galaxea fascicularis*, are known to produce copious amounts of mucous (Brown & Bythell, 2005). Major differences were observed in the expression of genes encoding mucins (Figure 3D), with many mucin-encoding genes more abundantly expressed in the STs. Mucins are a family of large glycoproteins (0.5–20 MDa) formed from a central protein core with heavily glycosylated side-chains, and are known to be key component in most gel-like secretions (Bansil & Turner, 2006). Mucins can be membrane-bound or secreted (gel-forming), and together, both types of mucin generate the characteristic mucus layer (Hollingsworth & Swanson, 2004; Thornton, Rousseau, & McGuckin, 2008). The eight genes more abundantly expressed in the ST were all homologs of MUC1, a membrane-bound mucin which is anchored directly to the cell membrane and is involved (in other organisms) in surface protection, cell-cell interactions, adhesion and signal transduction (Dhar & McAuley, 2019). The increase in mucocyte density and MUC1 gene expression in STs compared to CTs suggest that mucous may have a prominent role in the aggression mechanism, potentially as a “carrier” for toxins that are not derived from the stinging cells (Bartosz et al., 2008; Ben-Ari et al., 2018; Columbus-Shenkar et al., 2018; Moran et al., 2011).

In the CTs of corals, one of the roles of the mucous produced by the ectodermal cells is to produce a flow along the tentacles, bringing food particles towards the mount (Brown & Bythell, 2005). This process is powered by the movement of ectodermal cilia, and is called a ciliary-mucoid feeding. The lack of the cilia on the ectodermal surface of the STs supports the notion that these tentacles do not participate in feeding, and suggest that qualitatively different mucous might be involved in aggression compared to ciliary-mucoid feeding. In contrast, the endodermal cells of both tentacle types exhibited flagella, which are involved in circulating the gastrovascular fluid through the tentacles both as an internal transport system and as the driving force for the tentacles’ hydrostatic skeleton (Gladfelter, 1983). No clear differences were observed between the CTs and STs in the expression of genes involved in the synthesis of cilia and flagella (Supplementary excel Table). However, two histamine H2-like receptor genes were found more highly expressed in the CTs, and one such gene was more abundantly expressed in the body, with no such genes more abundantly expressed in the STs (Supplementary Excel Table). Histamine H2 receptors have been shown to play part in the mucociliary transport mechanisms in ascidians and mollusks, by controlling ciliary beat frequency (Cima & Franchi, 2016; P. Williams, Akande, Catapane, & Carroll, 2013). The lower expression of these genes in the ST may be due to the lack of cilia and of the muco-ciliary transport mechanism in these tentacles.

### Nematocyte development and regulation in the CT and the ST

Nematocytes are the main venom delivery system in cnidarians, and previous studies have shown that the nematocytes of the CTs and STs differ (Hidaka & Yamazato, 1984). Specifically, the acrosphere of the CTs contain mostly spirocysts and Microbasic p-Mastigophores (MpMs), whereas STs contain many Microbasic b-mastigophpores (MbMs) and somewhat fewer spirocysts (Hidaka & Yamazato, 1984). These differences were observed also between the CTs and STs in our animals, which were collected ~8,700 Km apart from those studied by Hidaka in 1984, and which may belong to different genotypes (Keshavmurthy et al., 2013). The three nematocyst types, MbM, MpM and HI (with the latter found primarily at the base of the tentacles), are all penetrating nematocytes, and thus may be used to deliver toxins into the target organism. The spirocysts, in contrast, do not penetrate the target, but rather entangle it. The higher abundance of spirocysts in the CTs may be related to the need to ensnare mobile prey organisms in order to feed on them, a requirement not shared by the STs.

In Hydra, nematocytes develop from their precursor cells in the body column, and during this period many of the proteins comprising the nematocyst structure are expressed, including nematogalectins and NOWA (Jung Shan Hwang et al., 2010). The mature nematocytes then migrate up the body column and into the tentacles, where they are mounted into the ectodermanl batteries. In contrast, little is known about the patterns of development of nematocysts in other cnidarians, although a NOWA-like gene is expressed in the tentacles of the primary polyp of *Nematostella* (Sunagar et al., 2018), and nematogalectins are expressed in the tentacle bulb of the jellyfish *Clytia hemisphaerica* where nematogenesis has been shown to occur (Denker, Manuel, Leclère, Le Guyader, & Rabet, 2008; Jung Shan Hwang et al., 2010). Our results showing the expression of nematogalectins and NOWA in the CTs and the ST, and the higher expression level in the STs compared to the CTs, suggest that the development of the nematocysts in *G. facsicularies*, including the change in nematocyte composition as the STs develop from CTs (Hidaka & Yamazato, 1984), occurs in the tentacles. We did not identify transcripts encoding Cnidoin or Spinalin, either in our transcriptome or in the recently published draft genome (Ying et al., 2018). This is in accordance with the suggestion that spinalin, which forms the spines decorating the tubules of some Hydra nematocysts, is restricted to *Hydra* (Milde et al., 2009), and highlights how little we know about the molecular underpinnings of the rich array of cnidarian nematocytes.

The discharge of nematocytes in cnidarians is a highly regulated process. While nematocytes may discharge when stimulated individually, they are innervated and regulated by the cnidarian nerve net (Tardent, 1995). Given that the nematocytes of the CTs need to discharge in response to mobile prey, whereas those in the STs need to respond to closely-related sessile targets, it is likely that the processes regulating their discharge are different. Little is known about the discharge mechanism of nematocytes involved in aggression, although an electrical “action potential” was measured in the acrorhagi of sea anemones in response to the identification of a non-self target (Lubbock & Shelton, 1981). Intriguingly, we identified two GO terms, voltage gates calcium and ionoptropic glutamate activity, as enriched in the STs compared with the CTs. Signaling by glutamate regulates nematocyte discharge in other cnidarians (Kass-Simon & Scappaticci, 2004). Similarly, in Hydra, light inhibits nematocyte discharge through opsin-mediated phototransduction (Plachetzki, Fong, & Oakley, 2012), and we identified non-visual photoreceptor (NVP) GPCR genes such as Melanopsin and Melatonin as more abundantly expressed in the STs. To what extent ionotropic glutamate transmission and photoreceptors are involved in mediating nematocyte discharge, or whether their role lies in mediating other behavioral aspects of the STs activity such as the search activity, remains to be determined.

### One organism, two tentacles, two functions, two venoms

The primary function of the catch and sweeper tentacles are to affect target organisms, presumably using nematocyst-derived venom or other, non-nematocystic, toxins. For the CTs, the targets are the coral’s prey or predators. In the case of prey, the venom needs to cause rapid paralysis, enabling the sessile predator to catch and eat mobile prey (Brodie, 2009). In the case of predator deterrence, pain production rather than paralysis might be a better ecological end-point (*e.g*. (Harris & Jenner, 2019; Inceoglu et al., 2003; Prince, Gunson, & Scarpa, 1985)). In contrast, the target of the ST are territorial competitors, and therefore the venom likely does not need to induce paralysis but rather to cause tissue damage (Bartosz et al., 2008). Consistent with this view, we observed both qualitative and quantitative differences in the toxicity profile between the CTs and the STs – only the CTss contain a paralytic (presumably neurotoxic) activity, whereas both tentacle types contain hemolytic and PLA2 activities, but the CT are more hemolytic and the STs have more PLA2 activities (Figure 4). This suggests that while the neurotoxic activity is dispensable for territorial aggression, the hemolytic and PLA2 activities may be relevant for both aggression and prey/predator encounter.

At the molecular level, different toxic proteins may be responsible for the observed hemolytic and PLA2 activities in each tentacle type. In the case of hemolysins, two out of four actinoporin transcripts were more abundantly expressed in the STs, whereas one actinoporin and a transcript encoding a CrTX-A like toxin were more abundantly expressed in the CTs. Actinoporins are widely distributed in cnidarians, including in corals (Ben-Ari et al., 2018), and in some cases have been shown to originate from the nematocysts (Basulto et al., 2006; J. S. Hwang et al., 2007; Rachamim et al., 2015). In addition, different nematocytes may each contain a different actinoporin (J. S. Hwang et al., 2007). In contrast, in the aggressive organs of the sea anemone *Actinia equina*, actinoporins are found in the tissue extract but not in the nematocyte venom (Bartosz et al., 2008), as shown for other toxins in *Nematostella vectensis* (Columbus-Shenkar et al., 2018; Moran et al., 2011). These observations suggest the potential for significant functional differences between different actinoporins. Importantly, pore-forming toxins, to which both the actinoporin and CrTX-A like toxins belong, can cause cell lysis but may also have neurotoxic-paralytic activities (Rohou, Nield, & Ushkaryov, 2007; Zhang, Fishman, Sher, & Zlotkin, 2003). We therefore speculate that the actinoporin and CrTX-A like toxins found in the CT may have stronger lethal activities, whereas those more abundantly expressed in the ST may induce tissue damage and potentially inflammation, potentially through the induction of cell death mechanisms in the target tissue (Ng et al., 2019; Soto et al., 2018). Importantly, while actinoporins have been intensively studied from evolutionary, biochemical and structural perspectives (*e.g*. as models of how water-soluble proteins interact with cell membranes, (Anderluh & Macek, 2003; Macrander & Daly, 2016)), little is known about their actual effect on the target organisms (prey, predators or competitors) or the ecological role of this protein family in cnidarians.

Similar to the hemolysins, different PLA2 genes were more abundantly expressed in each type of tentacle, again suggesting a potential functional differentiation between different toxins from the same family. We speculate that the PLA2 toxins more abundantly expressed in the CTs may have a neurotoxic effect (e.g. (Lotan, Fishman, & Zlotkin, 1996; Rigoni et al., 2005)), whereas the isoforms more abundantly expressed in the ST may cause tissue damage, potentially through the activation of the Lyso-PAF inflammation pathway in the target coral (Quinn et al., 2016).

Interestingly, PLA2 toxins, as well as metalloproteases and kunitz-type serine protease toxins, were all more abundantly expressed in the aggressive organs (acrorhagi) of the sea anemone *Anthopleura elegantissima* compared to non-aggressive polyps (Macrander, Brugler, & Daly, 2015). We also observed several metalloproteases and a putative Kunitz-type toxin that were more abundantly expressed in the ST (Figure 4A). Metalloproteases are important venom components of most venomous animals including cnidarians, where they can cause tissue degradation (Feitosa et al., 1998; Ponce, Brinkman, Potriquet, & Mulvenna, 2016; H. F. Williams et al., 2019). Metalloproteases, mostly belonging to the Astacin-like family, have previously been detected in the soluble nematocyst content of jellyfish, Hydra, sea anemones and corals (Gacesa et al., 2015; Jouiaei et al., 2015; R. Li et al., 2014; Moran et al., 2013; Rachamim et al., 2015). We speculate that these three toxin types - PLA2s, metalloproteases and Kunitz-type protease inhibitors - may form a “functional triad”, playing an important role in the tissue-degrading effect of venom in general, and the venom involved in territorial aggression in corals in particular.

## Summary and conclusions

The coral reef is a highly competitive place, where the morphologically simple and sessile corals need to be able to “fight” in parallel on different fronts – catching prey, defending themselves against predators and competing in territorial aggressive encounters with other corals. We propose a conceptual model describing some physiological and molecular features of the two different tentacle types of *Galaxea fascicularis*, each of which has evolved to address a different ecological challenge. The CTs are adapted to catch prey using paralytic toxins and hemolysins, likely injected through the MpM nematocytes, with spirocytes helping to entangle the prey. The prey is then transferred to the mouth using cilia combined with a mobile mucous layer, and the ciliary beating is controlled by the Histamine H2 system. The ST, in contrast, actively “search” for their long-distance targets, with the tentacle length and movement (and potentially the nematocyte discharge) controlled by the opsin and allostatin sensory systems. Once the target organism is identified, the ST deliver primarily enzymatic venom through the two different MbM nematocyte types, as well as (potentially) as part of the extensive non-nematocytic mucous secretion. We speculate that the ST mucous may have a higher relative composition of membrane-bound glycoproteins (such as mucins), thus being less of a “mobile” or flowing mucous cover. This mucous might also play a role in the recognition of self vs non-self tissue. This conceptual model is based on histological observations, functional toxicity assays and analyses of differential gene expression, and provides several hypotheses that can be tested using experimental tools currently available for non-model corals such as *Galaxea fascicularis*.

The venom of one organism – the coral – needs to produce three different functional outcomes – paralysis, pain and tissue degradation. The need for venom to fulfill different roles is not unique to corals - scorpions, cone snails and assassin bugs, for example, each produce multiple venom cocktails that nevertheless are injected through the same venom apparatus (Dutertre et al., 2014; Inceoglu et al., 2003; Walker et al., 2018). In the case of the coral, the morphological distinction between the CT and ST enable each to produce a unique chemical armament. However, it is also possible that within each tentacle type there may be more than one type of venom. For example, in Hydra, different nematocytes are discharged in response to prey and to predators (Ewer, 1947; D. Sher & Zlotkin, 2009). Additionally, many cnidarians also employ toxins that are not delivered through the stinging cells (Columbus-Shenkar et al., 2018; Moran et al., 2011; Daniel Sher et al., 2008). It remains to be tested whether, within each tentacle type, different effector molecules or toxins are delivered through different cell types. If so, this “division of labor” might enable a complex chemical armament to be delivered in a target-and time-specific manner.

Our results also leave many questions open. For example, the genes more abundantly expressed in the ST are enriched with the GO terms “oxygen binding” and “heme”, yet the relationship between these molecular functions and the biology of the STs is still unclear. Genes encoding RNAase, transcription factor binding and ubiquitin-protein transferase activities are also enriched in the ST, suggesting a constant dynamic remodeling of these tentacles, a process that has yet to be studied in detail. Finally, approximately 34% of the genes identified in the transcriptome could not be functionally annotated, including 4% of the genes differentially expressed between the CT and the ST. This highlights the richness of molecular functions still to be discovered that allow these corals to survive and thrive on the coral reef.

## Acknowledgements

This study was supported by grants 1239/13 and 312/15 from the Israel Science Foundation (to DS and TM, respectively) and by the associated grant number 2236/16 from the Israeli National Center for Personalized Medicine (to DS). We thank the Dive Team of the Underwater Observatory in Eilat for assistance with sample collection and the Crown Genomics institute of the Nancy and Stephen Grand Israel National Center for Personalized Medicine, Weizmann Institute of Science, for sequencing. Computations presented in this work were performed on the Hive computer cluster at the University of Haifa, which is partly funded by ISF grant 2155/15.

## Data accessibility statement

The raw sequencing reads, and the annotated Trinity assembly, are deposited with a temporary SRA project id SUB6897590 (we will provide a final accession when available).

## Author contributions

OY, TM and DS designed research, OY, YP, AM, MOL, TM and DS performed research and analyzed data, OY, TM and DS wrote the paper with contributions from all authors.

## References

Abelson, A., & Loya, Y. (1999). Interspecific aggression among stony corals in Eilat, red sea: A hierarchy of aggression ability and related parameters. Bulletin of Marine Science, 65(3), 851–860.

Alzugaray, M. E., Hernández-Martínez, S., & Ronderos, J. R. (2016). Somatostatin signaling system as an ancestral mechanism: Myoregulatory activity of an Allatostatin-C peptide in Hydra. Peptides, 82, 67–75. doi:https://doi.org/10.1016/j.peptides.2016.05.011

Anderluh, G., & Macek, P. (2003). Dissecting the actinoporin pore-forming mechanism. Structure (Camb), 11(11), 1312–1313.

Atrigenio, M., Ali, #xf1, o, P., & Conaco, C. (2017). Influence of the Blue Coral Heliopora coerulea on Scleractinian Coral Larval Recruitment. Journal of Marine Biology, 2017, 5. doi:10.1155/2017/6015143

Bansil, R., & Turner, B. S. (2006). Mucin structure, aggregation, physiological functions and biomedical applications. Current Opinion in Colloid & Interface Science, 11(2), 164–170. doi:https://doi.org/10.1016/j.cocis.2005.11.001

Bartosz, G., Finkelshtein, A., Przygodzki, T., Bsor, T., Nesher, N., Sher, D., & Zlotkin, E. (2008). A pharmacological solution for a conspecific conflict: ROS-mediated territorial aggression in sea anemones. Toxicon, 51(6), 1038–1050.

Basulto, A., Perez, V. M., Noa, Y., Varela, C., Otero, A. J., & Pico, M. C. (2006). Immunohistochemical targeting of sea anemone cytolysins on tentacles, mesenteric filaments and isolated nematocysts of Stichodactyla helianthus. Journal of Experimental Zoology Part a-Comparative Experimental Biology, 305A(3), 253–258.

Beckmann, A., & Ozbek, S. (2012). The nematocyst: a molecular map of the cnidarian stinging organelle. International Journal of Developmental Biology, 56(6-8), 577–582. doi:113472ab [pii] 10.1387/ijdb.113472ab

Ben-Ari, H., Paz, M., & Sher, D. (2018). The chemical armament of reef-building corals: inter-and intra-specific variation and the identification of an unusual actinoporin in Stylophora pistilata. Scientific Reports, 8(1), 251. doi:10.1038/s41598-017-18355-1

Bengtsson-Palme, J., Hartmann, M., Eriksson, K. M., Pal, C., Thorell, K., Larsson, D. G. J., & Nilsson, R. H. (2015). metaxa2: improved identification and taxonomic classification of small and large subunit rRNA in metagenomic data. Molecular Ecology Resources, 15(6), 1403–1414. doi:10.1111/1755-0998.12399

Bolger, A. M., Lohse, M., & Usadel, B. (2014). Trimmomatic: a flexible trimmer for Illumina sequence data. Bioinformatics, 30(15), 2114–2120. doi:10.1093/bioinformatics/btu170

Brodie, E. D., III. (2009). Toxins and venoms. Current Biology, 19(20), R931–R935. doi:10.1016/j.cub.2009.08.011

Brown, B. E., & Bythell, J. C. (2005). Perspectives on mucus secretion in reef corals. Mar Ecol Prog Ser, 296.

Casewell, N. R. (2012). On the ancestral recruitment of metalloproteinases into the venom of snakes. Toxicon, 60(4), 449–454. doi:https://doi.org/10.1016/j.toxicon.2012.02.006

Chadwick, N. E. (1988). Competition and locomotion in a free-living fungiid coral. Journal of Experimental Marine Biology and Ecology, 123(3), 189–200. doi:https://doi.org/10.1016/0022-0981(88)90041-X

Chadwick, N. E., & Morrow, K. M. (2011). Competition Among Sessile Organisms on Coral Reefs. In Z. Dubinsky & N. Stambler (Eds.), Coral Reefs: An Ecosystem in Transition (pp. 347–371). Dordrecht: Springer Netherlands.

Chornesky, E. A. (1983). Induced Development of Sweeper Tentacles on the Reef Coral Agaricia agaricites: A Response to Direct Competition. Biological Bulletin, 165(3), 569–581. doi:10.2307/1541466

Cima, F., & Franchi, N. (2016). Histamine Stimulates Ciliary Beat Frequency via the H2 Receptor in the Protochordate Botryllus schlosseri. Journal of Experimental Zoology Part B: Molecular and Developmental Evolution, 326(3), 176–192. doi:10.1002/jez.b.22675

Columbus-Shenkar, Y. Y., Sachkova, M. Y., Macrander, J., Fridrich, A., Modepalli, V., Reitzel, A. M., … Moran, Y. (2018). Dynamics of venom composition across a complex life cycle. eLife, 7, e35014. doi:10.7554/eLife.35014

Conesa, A., Gotz, S., Garcia-Gomez, J. M., Terol, J., Talon, M., & Robles, M. (2005). Blast2GO: a universal tool for annotation, visualization and analysis in functional genomics research. Bioinformatics, 21(18), 3674–3676. doi:bti610 [pii] 10.1093/bioinformatics/bti610

Connell, J. H., Hughes, T. P., Wallace, C. C., Tanner, J. E., Harms, K. E., & Kerr, A. M. (2004). A LONG-TERM STUDY OF COMPETITION AND DIVERSITY OF CORALS. Ecological Monographs, 74(2), 179–210. doi:10.1890/02-4043

Denker, E., Manuel, M., Leclère, L., Le Guyader, H., & Rabet, N. (2008). Ordered progression of nematogenesis from stem cells through differentiation stages in the tentacle bulb of Clytia hemisphaerica (Hydrozoa, Cnidaria). Developmental Biology, 315(1), 99–113. doi:http://dx.doi.org/10.1016/j.ydbio.2007.12.023

Dhar, P., & McAuley, J. (2019). The Role of the Cell Surface Mucin MUC1 as a Barrier to Infection and Regulator of Inflammation. Frontiers in cellular and infection microbiology, 9, 117–117. doi:10.3389/fcimb.2019.00117

Dutertre, S., Jin, A.-H., Vetter, I., Hamilton, B., Sunagar, K., Lavergne, V., … Lewis, R. J. (2014). Evolution of separate predation-and defence-evoked venoms in carnivorous cone snails. Nature Communications, 5(1), 3521. doi:10.1038/ncomms4521

Einat, D. L., & Nanette, E. C. (2006). Long-term effects of competition on coral growth and sweeper tentacle development. Marine Ecology Progress Series, 313, 115–123.

Ertman, S. C., & Davenport, D. (1981). Tentacular Nematocyte Discharge and “Self-Recognition” in Anthopleura elegantissima Brandt. Biological Bulletin, 161(3), 366–370. doi:10.2307/1540941

Evensen, N. R., & Edmunds, P. J. (2018). Effect of elevated pCO2 on competition between the scleractinian corals Galaxea fascicularis and Acropora hyacinthus. Journal of Experimental Marine Biology and Ecology, 500, 12–17. doi:https://doi.org/10.1016/j.jembe.2017.12.002

Ewer, R. F. (1947). On the functions and mode of action of the nematocysts of Hydra. Proc. Zool. Soc. Lond, 117, 365–376.

Fearon, R. J., & Cameron, A. M. (1997). Preliminary Evidence Supporting the Ability of Hermatypic Corals to Affect Adversely Larvae and Early Settlement Stages of Hard Coral Competitors. Journal of Chemical Ecology, 23(7), 1769–1780. doi:10.1023/B:JOEC.0000006450.55638.b2

Feitosa, L., Gremski, W., Veiga, S. S., Elias, M. C. Q. B., Graner, E., Mangili, O. C., & Brentani, R. R. (1998). Detection and characterization of metalloproteinases with gelatinolytic, fibronectinolytic and fibrinogenolytic activities in Brown spider (Loxosceles intermedia) venom. Toxicon, 36(7), 1039–1051. doi:https://doi.org/10.1016/S0041-0101(97)00083-4

Gacesa, R., Chung, R., Dunn, S. R., Weston, A. J., Jaimes-Becerra, A., Marques, A. C., … Long, P. F. (2015). Gene duplications are extensive and contribute significantly to the toxic proteome of nematocysts isolated from Acropora digitifera (Cnidaria: Anthozoa: Scleractinia). BMC Genomics, 16(1), 774. doi:10.1186/s12864-015-1976-4

Genin, A., Karp, L., & Miroz, A. (1994). Effects of Flow on Competitive Superiority in Scleractinian Corals. Limnology and Oceanography, 39(4), 913–924.

Gladfelter, E. H. (1983). Circulation of fluids in the gastrovascular system of the reef coral Acropora cervicornis. The Biological Bulletin, 165(3), 619–636. doi:10.2307/1541469

Goldberg, W. M., Grange, K. R., Taylor, G. T., & Zuniga, A. L. (1990). The Structure of Sweeper Tentacles in the Black Coral Antipathes fiordensis. The Biological Bulletin, 179(1), 96–104. doi:10.2307/1541743

Grabherr, M. G., Haas, B. J., Yassour, M., Levin, J. Z., Thompson, D. A., Amit, I., … Regev, A. (2011). Full-length transcriptome assembly from RNA-Seq data without a reference genome. Nature Biotechnology, 29(7), 644–652. doi:10.1038/nbt.1883nbt.1883 [pii]

Gundlach, K. A., & Watson, G. M. (2018). Self/Non-Self Recognition Affects Cnida Discharge and Tentacle Contraction in the Sea Anemone Haliplanella luciae. The Biological Bulletin, 235(2), 83–90. doi:10.1086/699564

Hall, M. R., Kocot, K. M., Baughman, K. W., Fernandez-Valverde, S. L., Gauthier, M. E. A., Hatleberg, W. L., … Degnan, B. M. (2017). The crown-of-thorns starfish genome as a guide for biocontrol of this coral reef pest. Nature, 544(7649), 231–234. doi:10.1038/nature22033

Harris, R. J., & Jenner, R. A. (2019). Evolutionary Ecology of Fish Venom: Adaptations and Consequences of Evolving a Venom System. Toxins, 11(2), 60. doi:10.3390/toxins11020060

Hidaka, M. (1985). Nematocyst Discharge, Histoincompatibility, and the Formation of Sweeper Tentacles in the Coral Galaxea fascicularis. Biological Bulletin, 168(3), 350–358. doi:10.2307/1541517

Hidaka, M., & Yamazato, K. (1984). Intraspecific interactions in a scleractinian coral, Galaxea fascicularis: Induced formation of sweeper tentacles. Coral Reefs, 3(2), 77–85. doi:10.1007/bf00263757

Hollingsworth, M. A., & Swanson, B. J. (2004). Mucins in cancer: protection and control of the cell surface. Nature Reviews Cancer, 4(1), 45–60. doi:10.1038/nrc1251

Hwang, J. S., Ohyanagi, H., Hayakawa, S., Osato, N., Nishimiya-Fujisawa, C., Ikeo, K., … Gojobori, T. (2007). The evolutionary emergence of cell type-specific genes inferred from the gene expression analysis of Hydra. Proceedings of the National Academy of Sciences of the United States of America, 104(37), 14735–14740. doi:0703331104 [pii] 10.1073/pnas.0703331104

Hwang, J. S., Takaku, Y., Momose, T., Adamczyk, P., Özbek, S., Ikeo, K., … Gojobori, T. (2010). Nematogalectin, a nematocyst protein with GlyXY and galectin domains, demonstrates nematocyte-specific alternative splicing in Hydra. Proceedings of the National Academy of Sciences, 107(43), 18539–18544. doi:10.1073/pnas.1003256107

Inceoglu, B., Lango, J., Jing, J., Chen, L., Doymaz, F., Pessah, I. N., & Hammock, B. D. (2003). One scorpion, two venoms: prevenom of Parabuthus transvaalicus acts as an alternative type of venom with distinct mechanism of action. Proceedings of the National Academy of Sciences of the United States of America, 100(3), 922–927.

Jackson, J. B., & Buss, L. (1975). Alleopathy and spatial competition among coral reef invertebrates. Proceedings of the National Academy of Sciences of the United States of America, 72(12), 5160–5163. doi:10.1073/pnas.72.12.5160

Jouiaei, M., Sunagar, K., Federman Gross, A., Scheib, H., Alewood, P. F., Moran, Y., & Fry, B. G. (2015). Evolution of an Ancient Venom: Recognition of a Novel Family of Cnidarian Toxins and the Common Evolutionary Origin of Sodium and Potassium Neurotoxins in Sea Anemone. Molecular Biology and Evolution, 32(6), 1598–1610. doi:10.1093/molbev/msv050

Kass-Simon, G., & Scappaticci, A. A. (2004). Glutamatergic and GABAnergic control in the tentacle effector systems of Hydra vulgaris, Dordrecht.

Keshavmurthy, S., Yang, S.-Y., Alamaru, A., Chuang, Y.-Y., Pichon, M., Obura, D., … Chen, C. A. (2013). DNA barcoding reveals the coral “laboratory-rat”, Stylophora pistillata encompasses multiple identities. Scientific Reports, 3, 1520. doi:10.1038/srep01520

Lang, J. (1973). Interspecific aggression by scleractinian corals. 2. Why the race is not only to the swift. Bulletin of Marine Science, 23, 260–179.

Lang, J., & Chornesky, E. A. (1990). Competition between scleratinian reef corals - a review of mechanisms and effects. In Z. Dubinsky (Ed.), Ecosystems of the world. V. 25. Coral reefs (pp. 209–257): Elsevier.

Lee, H., Jung, E.-s., Kang, C., Yoon, W. D., Kim, J.-S., & Kim, E. (2011). Scyphozoan jellyfish venom metalloproteinases and their role in the cytotoxicity. Toxicon, 58(3), 277–284. doi:https://doi.org/10.1016/j.toxicon.2011.06.007

Li, B., & Dewey, C. N. (2011). RSEM: accurate transcript quantification from RNA-Seq data with or without a reference genome. BMC Bioinformatics, 12(1), 323. doi:10.1186/1471-2105-12-323

Li, R., Yu, H., Xue, W., Yue, Y., Liu, S., Xing, R., & Li, P. (2014). Jellyfish venomics and venom gland transcriptomics analysis of Stomolophus meleagris to reveal the toxins associated with sting. Journal of Proteomics, 106, 17–29. doi:https://doi.org/10.1016/j.jprot.2014.04.011

Lotan, A., Fishman, L., & Zlotkin, E. (1996). Toxin compartmentation and delivery in the cnidaria: The nematocyst’s tubule as a ultiheaded poisonous arrow. The journal of Experimental Zoology, 275, 444–451.

Love, M. I., Huber, W., & Anders, S. (2014). Moderated estimation of fold change and dispersion for RNA-seq data with DESeq2. Genome Biology, 15(12), 550. doi:10.1186/s13059-014-0550-8

Lubbock, R., & Shelton, G. A. B. (1981). Electrical activity following cellular recognition of self and non-self in a sea anemone. Nature, 289(1), 59.

Macrander, J., Brugler, M. R., & Daly, M. (2015). A RNA-seq approach to identify putative toxins from acrorhagi in aggressive and non-aggressive Anthopleura elegantissima polyps. BMC Genomics, 16(1), 221. doi:10.1186/s12864-015-1417-4

Macrander, J., & Daly, M. (2016). Evolution of the Cytolytic Pore-Forming Proteins (Actinoporins) in Sea Anemones. Toxins, 8(12), 368.

Milde, S., Hemmrich, G., Anton-Erxleben, F., Khalturin, K., Wittlieb, J., & Bosch, T. C. G. (2009). Characterization of taxonomically restricted genes in a phylum-restricted cell type. Genome Biology, 10(1). doi:Artn R8 Doi 10.1186/Gb-2009-10-1-R8

Moran, Y., Genikhovich, G., Gordon, D., Wienkoop, S., Zenkert, C., Özbek, S., … Gurevitz, M. (2011). Neurotoxin localization to ectodermal gland cells uncovers an alternative mechanism of venom delivery in sea anemones. Proceedings of the Royal Society B: Biological Sciences. doi:10.1098/rspb.2011.1731

Moran, Y., Praher, D., Schlesinger, A., Ayalon, A., Tal, Y., & Technau, U. (2013). Analysis of soluble protein contents from the nematocysts of a model sea anemone sheds light on venom evolution. Marine biotechnology (New York, N.Y.), 15(3), 329–339. doi:10.1007/s10126-012-9491-y

Mullen, A. D., Treibitz, T., Roberts, P. L. D., Kelly, E. L. A., Horwitz, R., Smith, J. E., & Jaffe, J. S. (2016). Underwater microscopy for in situ studies of benthic ecosystems. Nature Communications, 7(1), 12093. doi:10.1038/ncomms12093

Nagai, H., Takuwa, K., Nakao, M., Ito, E., Masami, M., Noda, M., & Nakajima, T. (2000). Novel proteinaceous toxins from the Box Jellyfish (Sea Wasp) Carybdea rastoni. Biochemical and Biophysical Research Communications, 275, 582–588.

Ng, T. J., Teo, M. Y. M., Liew, D. S., Effiong, P. E., Hwang, J. S., Lim, C. S. Y., & In, L. L. A. (2019). Cytotoxic and apoptosis-inducing effects of wildtype and mutated Hydra actinoporin-like toxin 1 (HALT-1) on various cancer cell lines. PeerJ, 7, e6639. doi:10.7717/peerj.6639

Peach, M. B., & Hoegh-Guldberg, O. (1999). Sweeper Polyps of the Coral Goniopora tenuidens (Scleractinia: Poritidae). Invertebrate Biology, 118(1), 1–7. doi:10.2307/3226906

Plachetzki, D. C., Fong, C. R., & Oakley, T. H. (2012). Cnidocyte discharge is regulated by light and opsin-mediated phototransduction. BMC Biology, 10(1), 17. doi:10.1186/1741-7007-10-17

Ponce, D., Brinkman, D. L., Potriquet, J., & Mulvenna, J. (2016). Tentacle Transcriptome and Venom Proteome of the Pacific Sea Nettle, Chrysaora fuscescens (Cnidaria: Scyphozoa). Toxins, 8(4), 102.

Prince, R. C., Gunson, D. E., & Scarpa, A. (1985). Sting like a bee! The ionophoric properties of melittin. Trends in Biochemical Sciences, 10, 99.

Pritchard, L., & Dufton, M. J. (1999). Evolutionary trace analysis of the Kunitz/BPTI family of proteins: functional divergence may have been based on conformational adjustment. Journal of Molecular Biology, 285(4), 1589–1607.

Quinn, R. A., Vermeij, M. J. A., Hartmann, A. C., d’Auriac, I. G., Benler, S., Haas, A., … Rohwer, F. (2016). Metabolomics of reef benthic interactions reveals a bioactive lipid involved in coral defence. Proceedings of the Royal Society B: Biological Sciences, 283(1829), 20160469. doi:doi:10.1098/rspb.2016.0469

Rachamim, T., Morgenstern, D., Aharonovich, D., Brekhman, V., Lotan, T., & Sher, D. (2015). The dynamically evolving nematocyst content of an anthozoan, a scyphozoan, and a hydrozoan. Molecular Biology and Evolution, 32. doi:10.1093/molbev/msu335

Reed, L. J., & Muench, S. (1938). A simple method of estimating fifty per cent end point. Am. J. Hyg., 27, 493.

Rigoni, M., Caccin, P., Gschmeissner, S., Koster, G., Postle, A. D., Rossetto, O., … Montecucco, C. (2005). Equivalent Effects of Snake PLA2 Neurotoxins and Lysophospholipid-Fatty Acid Mixtures. Science, 310(5754), 1678–1680. doi:10.1126/science.1120640

Rohou, A., Nield, J., & Ushkaryov, Y. A. (2007). Insecticidal toxins from black widow spider venom. Toxicon : official journal of the International Society on Toxinology, 49(4), 531–549. doi:10.1016/j.toxicon.2006.11.021

Schmidt, C. A., Daly, N. L., & Wilson, D. T. (2019). Coral Venom Toxins. Frontiers in Ecology and Evolution, 7(320). doi:10.3389/fevo.2019.00320

Sebens, K. P., & Miles, J. S. (1988). Sweeper Tentacles in a Gorgonian Octocoral: Morphological Modifications for Interference Competition. Biological Bulletin, 175(3), 378–387. doi:10.2307/1541729

Seneca, F. O., & Palumbi, S. R. (2015). The role of transcriptome resilience in resistance of corals to bleaching. Molecular Ecology, 24(7), 1467–1484. doi:10.1111/mec.13125

Sher, D., Fishman, Y., Melamed-Book, N., Zhang, M., & Zlotkin, E. (2008). Osmotically driven prey disintegration in the gastrovascular cavity of the green hydra by a pore-forming protein. FASEB Journal, 22 (1), 207–221.

Sher, D., & Zlotkin, E. (2009). A hydra with many heads: protein and polypeptide toxins from hydra and their biological roles. Toxicon, 54(8), 1148–1161. doi:S0041-0101(09)00144-5 [pii] 10.1016/j.toxicon.2009.02.036

Soto, C., Bergado, G., Blanco, R., Griñán, T., Rodríguez, H., Ros, U., … Álvarez, C. (2018). Sticholysin II-mediated cytotoxicity involves the activation of regulated intracellular responses that anticipates cell death. Biochimie, 148, 18–35. doi:https://doi.org/10.1016/j.biochi.2018.02.006

Stabili, L., Schirosi, R., Parisi, M. G., Piraino, S., & Cammarata, M. (2015). The Mucus of Actinia equina (Anthozoa, Cnidaria): An Unexplored Resource for Potential Applicative Purposes. Marine Drugs, 13(8), 5276–5296.

Sunagar, K., Columbus-Shenkar, Y. Y., Fridrich, A., Gutkovich, N., Aharoni, R., & Moran, Y. (2018). Cell type-specific expression profiling unravels the development and evolution of stinging cells in sea anemone. BMC Biology, 16(1), 108. doi:10.1186/s12915-018-0578-4

Tardent, P. (1995). The cnidarian cnidocyte, a high-tech cellular weaponry. Bioessays, 17(4), 351–362.

Thornton, D. J., Rousseau, K., & McGuckin, M. A. (2008). Structure and Function of the Polymeric Mucins in Airways Mucus. Annual Review of Physiology, 70(1), 459–486. doi:10.1146/annurev.physiol.70.113006.100702

Walker, A. A., Mayhew, M. L., Jin, J., Herzig, V., Undheim, E. A. B., Sombke, A., … King, G. F. (2018). The assassin bug Pristhesancus plagipennis produces two distinct venoms in separate gland lumens. Nature Communications, 9(1), 755. doi:10.1038/s41467-018-03091-5

Williams, H. F., Mellows, B. A., Mitchell, R., Sfyri, P., Layfield, H. J., Salamah, M., … Vaiyapuri, S. (2019). Mechanisms underpinning the permanent muscle damage induced by snake venom metalloprotease. PLoS Neglected Tropical Diseases, 13(1), e0007041. doi:10.1371/journal.pntd.0007041

Williams, P., Akande, P., Catapane, E. J., & Carroll, M. A. (2013). Further studies on the sensory motor integration of gill lateral cilia in the bivalve mollusc Crassostrea virginica. The FASEB Journal, 27(1_supplement), 1120.1121–1120.1121. doi:10.1096/fasebj.27.1_supplement.1120.1

Williams, R. B. (1991). Acrorhagi, catch tentacles and sweeper tentacles: a synopsis of ‘aggression’ of actiniarian and scleractinian Cnidaria. Hydrobiologia, 216(1), 539–545. doi:10.1007/bf00026511

Ying, H., Cooke, I., Sprungala, S., Wang, W., Hayward, D. C., Tang, Y., … Miller, D. J. (2018). Comparative genomics reveals the distinct evolutionary trajectories of the robust and complex coral lineages. Genome Biology, 19(1), 175. doi:10.1186/s13059-018-1552-8

Zhang, M., Fishman, Y., Sher, D., & Zlotkin, E. (2003). Hydralysin, a novel animal group-selective paralytic and cytolytic protein from a noncnidocystic origin in hydra. Biochemistry, 42(30), 8939–8944.

Zlotkin, E., Fraenkel, G., Miranda, F., & Lissittzky, S. (1971). The effect of scorpion venom on blowfly larvae; a new method for the evaluation of scorpion venom potency. Toxicon, 9, 1–8.

